# Spaceflight decelerates the epigenetic clock orchestrated with a global alteration in DNA methylome and transcriptome in the mouse retina

**DOI:** 10.1101/2020.10.23.352658

**Authors:** Zhong Chen, Seta Stanbouly, Nina C. Nishiyama, Xin Chen, Michael D. Delp, Hongyu Qiu, Xiao W. Mao, Charles Wang

## Abstract

Astronauts exhibit an assortment of clinical abnormalities in their eyes during long-duration spaceflight. The purpose of this study was to determine whether spaceflight induces epigenomic and transcriptomic reprogramming in the retina or alters the epigenetic clock. The mice were flown for 37 days in animal enclosure modules on the International Space Station; ground-based control animals were maintained under similar housing conditions. Mouse retinas were isolated and both DNA methylome and transcriptome were determined by deep sequencing. We found that a large number of genes were differentially methylated with spaceflight, whereas there were fewer differentially expressed genes at the transcriptome level. Several biological pathways involved in retinal diseases such as macular degeneration were significantly altered. Our results indicated that spaceflight decelerated the retinal epigenetic clock. This study demonstrates that spaceflight impacts the retina at the epigenomic and transcriptomic levels, and such changes could be involved in the etiology of eye-related disorders among astronauts.

## Introduction

The spaceflight environment contains two major risk factors, irradiation and microgravity, which are thought to induce adverse effects on astronauts’ vision and central nervous systems^1,2,3^, and in particular, the eye and retina^4,5^. Studies reported that more than 30% of astronauts returning from Space Shuttle missions or the International Space Station (ISS) were diagnosed with eye problems that reduced visual acuity^3,6^. Furthermore, a recent study showed that spaceflight induced significant apoptosis in the retinal inner nuclear layer (INL) and ganglion cell layer (GCL) of mice^7^. Previous work has also shown that many genes involved in oxidative stress and mitochondria-associated apoptosis are altered in the retinas of space flown mice, and oxidative stress-induced apoptosis plays an important role in the response to space irradiation in mouse retinas^8,9,10^.

Previous research designed to better understand space-induced visual impairment in astronauts has primarily utilized proteomics and microarray technologies to better understand the biological effects of spaceflight on the mouse retina; these studies collectively reveal that spaceflight alters the expression of certain genes and proteins involved in cell death, mitochondrial function, endoplasmic reticulum (ER) stress, neuronal and glial cell loss, and axonal degeneration in mice^8,11^. Although there are studies analyzing the transcriptome in human retinas, none have been done under spaceflight conditions^12,13,14,15.^ Thus, the mechanisms underlying the spaceflight induced vision impairment remain largely unknown. Furthermore, the spaceflight-induced epigenomic and transcriptomic reprogramming in the retina has not yet been reported. In particularly, there has been no study of the effects of spaceflight on the epigenetic clocks of animals or humans.

There is great enthusiasm both within the scientific community and society in general with the ambitious plans for the return of manned missions to the moon and long-duration (>300 days) space travel to Mars and beyond. However, a comprehensive assessment of the adverse effects on the human body is still incomplete. In a recent NASA Twins Study, some persistent changes in gene expression, DNA damage, telomeres, neuro-ocular and cognitive function were observed when comparing the spaceflight twin brother to his earth-bound twin^16^. It has also been shown that long-duration spaceflight changes gene regulation at both the transcriptional and epigenetic levels, although the majority of these changes return to the preflight level six months after returning to earth. As part of a NASA research consortium, we investigated the DNA methylome, epigenetic clock and transcriptome of retinas from mice flown on a long-duration 37-day mission, the human equivalent of 4.1 years, as part of the Rodent Research-1 (RR-1) project. We identified differentially methylated genes (DMGs) and differentially expressed genes (DEGs) between spaceflight and ground control mice. In addition, as epigenetic clocks have been used to predict tissue biological ages and age-related diseases in both humans^17^ and mice^18,19^, we calculated spaceflight-induced epigenetic clock based on the DNA methylation at unique CpG sites of mouse retinas to predict the retinal tissue aging due to the spaceflight. This study provides new insights into the visual impairment due to long-duration spaceflight.

## Results

### 1. Study design and overall QC of RRBS DNA methylome and RNA-seq transcriptome data

To study the effects of spaceflight on retinal tissue epigenomic and transcriptomic changes, we constructed 18 reduced representation bisulfite sequencing (RRBS) libraries and one sample was removed from downstream analysis due to low quality (8 flight vs. 9 ground control mice) and 18 RNA-seq libraries (9 flight vs. 9 ground control mice). **Figure 1** shows the study design in detail. Fragment sizes of RRBS library were between 200 and 500 bp, with a peak around 300 bp. RNA-seq library size distribution ranged between 200-500 bp, with a peak around 280 bp. To monitor the quality of the RNA-seq, we also added External RNA Control Consortium (ERCC) spike-in controls in an amount equivalent to about 1% of the total RNA in each sample before RNA-seq library construction. We generated ~778 million reads of 75-bp single-end RRBS DNA methylome data and 764 million reads (75 bp X 2) of pair-end RNA-seq transcriptomic data, corresponding to an average of ~45.8 million (N=17) sequence reads per RRBS sample, and 42.5 million (N=18) sequence reads per RNA-seq sample.

**Figure 1.**
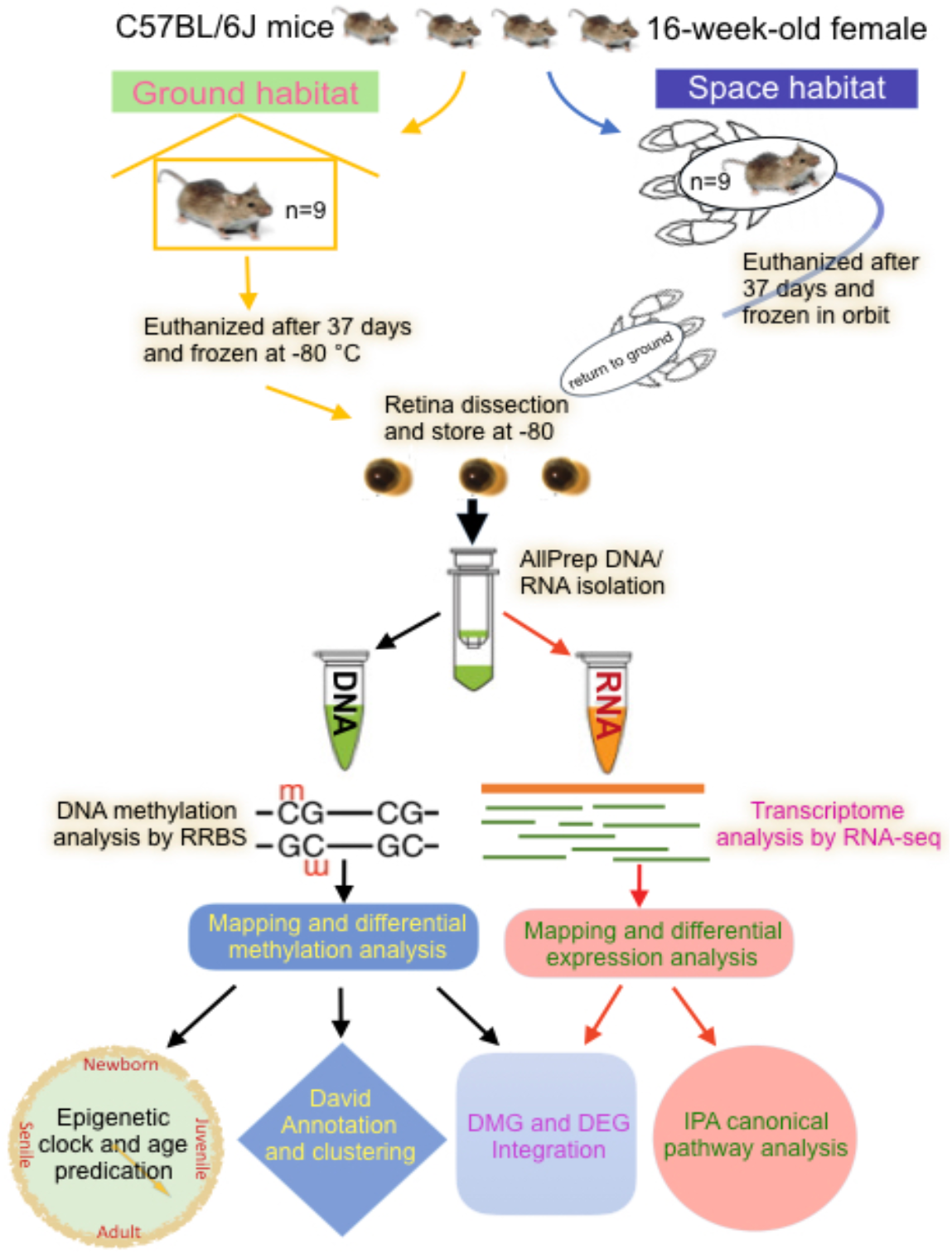
Study design. This study was under the NASA Rodent Research-1 (RR-1) project consortium. Spaceflight mice after flight were euthanized and frozen on orbit. The retinas were isolated from both the ground habitat control and flight mice after returning to ground. RNA and DNA were extracted from the retinas, and then RRBS and RNA-seq libraries were constructed to obtain DNA methylome and transcriptome, respectively. Retinal epigenetic age was calculated using MouseEpigeneticClock script. Disease and bio-functional pathways were generated using the DAVID GO functional annotation and the Ingenuity Pathway through Analysis (IPA) based on DMGs & DEGs.

High quality reads were obtained from both RRBS and RNA-seq data (**Supplementary Figure 1**). We aligned the RRBS DNA-seq methylome data, 75 bp single reads to the NCBI mouse GRCm38 genome. On average, 57.39% of the reads were aligned to the genome with an average of 26.3 million (N=17) aligned reads per sample. The aligned reads were further annotated, resulting in an average of 1.84 million (N=17) CpG sites covered by at least 10 reads. For RNA-seq transcriptomic data, we aligned the 75×2 bp fastq reads to the NCBI mouse GRCm38 genome, NCBI GRCm38 gene model, and ERCC transcripts. Overall, ~89.13% (N=18) of the reads were aligned to the genome, and 0.35% (N=18) to the ERCC transcripts. Among the reads aligned to the genome, 56.03% (N=18) to the exons, 20.88% (N=18) to the intron. Scatterplots of ERCC log2 (FPKM) vs. log2 (spike-in concentration) showed an overall linear relationship (R^2^ = 0.94) between the observed and the expected transcripts of the ERCC spike-in controls (**Supplementary Figure 1g**).

### 2. Global epigenomic and transcriptomic changes in the retinas of spaceflight mice

Overall, 31.2 million CpGs, ranging from 1.6 million to 2.1 million CpGs per sample, were used to study the genome-wide CpG methylation patterns in terms of the relationship to the genomic features. The profiling CpG methylation patterns revealed that the spaceflight mice were hypomethylated globally compared to ground mice (p=0.034, two-tailed t-test, **Figure 2a**). A significantly lower CpG methylation level was observed within CG islands compared to CpG shores in both ground and spaceflight mice (p < 2.2e^−16^, two-tailed t-test). In addition, the level of methylation was significantly higher in the introns compared to exons in both groups (p < 2.2e^−16^, two-tailed t-test). The methylation at the transcription start site (TSS) had the lowest methylation compared to other genomic regions (promoter, exon and intron) in both spaceflight and ground mice. Furthermore, 62.5% of the 643 differentially methylated genes (DMGs) were hypomethylated and 37.5% were hypermethylated (**Figure 2c**). Annotation of differentially methylated CpGs (DMCs) revealed that a large portion of the DMCs were located in introns, followed by intergenic regions and exons. About 15% of the DMCs were located in promoters (**Figure 2d**). Principal component analysis (PCA) (**Figure 3a**) and methylation heat map on DMCs showed a clear difference between spaceflight and ground control mice (**Figure 3c**). Overall, the methylation data clearly demonstrated that spaceflight induced profound DNA methylation changes.

**Figure 2.**
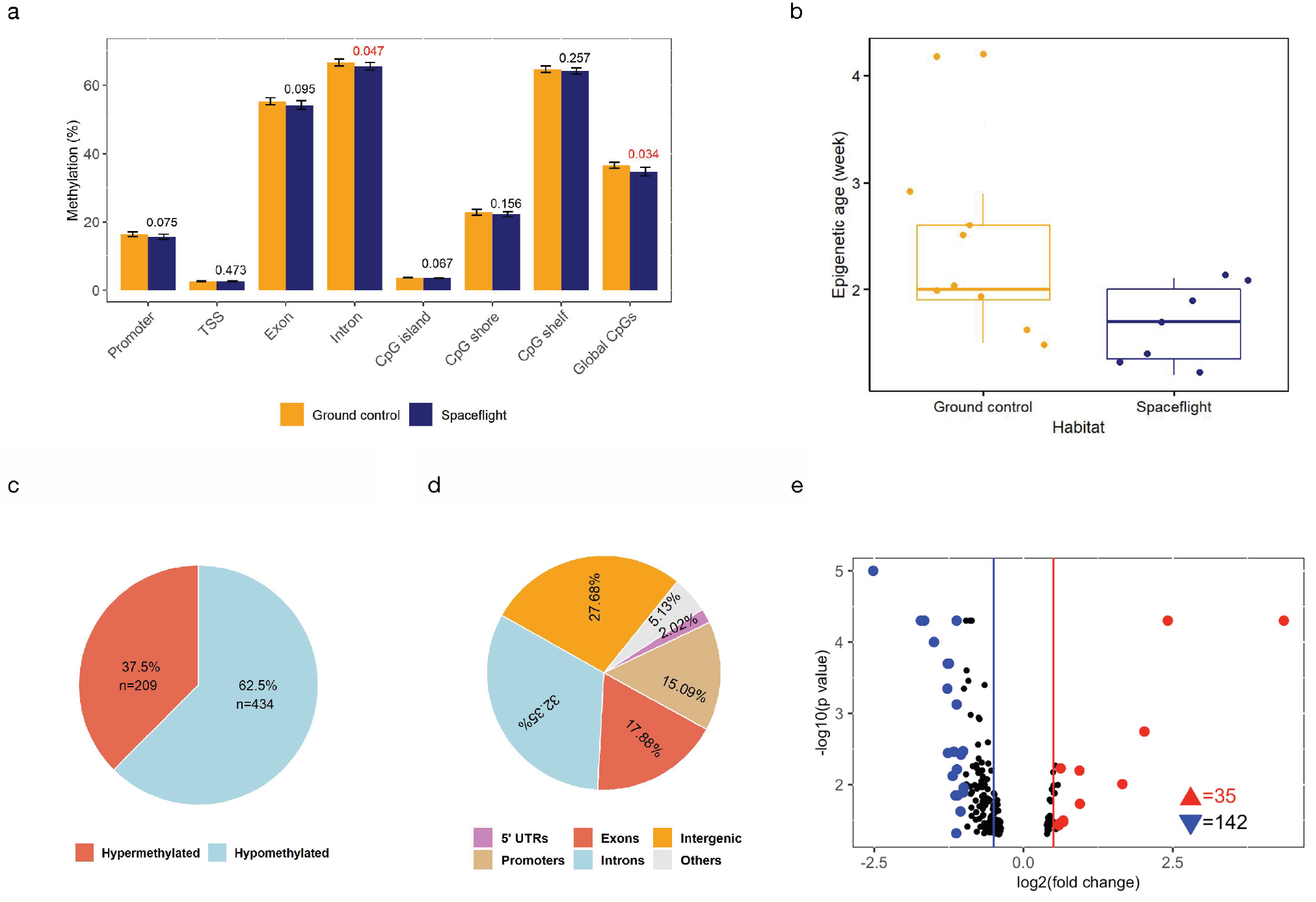
Spaceflight caused global epigenomic and transcriptomic changes in mice retinas. (**a**) Genome-wide CpG methylation status at various genome regions; (**b**) Epigenetic ages calculated using Stubb’s age estimator based on retinal methylomes (note that the chronological age at the euthanizing time was 21.2 weeks); (**c, d**) Characteristics of significant differentially methylated CpGs (DMCs, methylation change >10% and P ≤ 0.05); (**e**) Volcano plot showing significant differentially expressed genes (DEGs, fold change > 1.2 and P ≤ 0.05). Red and blue triangle indicates the number of up-regulated or down-regulated DEGs, respectively. DEGs with more than 50% expression change are highlighted in red (up-regulated) and blue (down-regulated).

**Figure 3.**
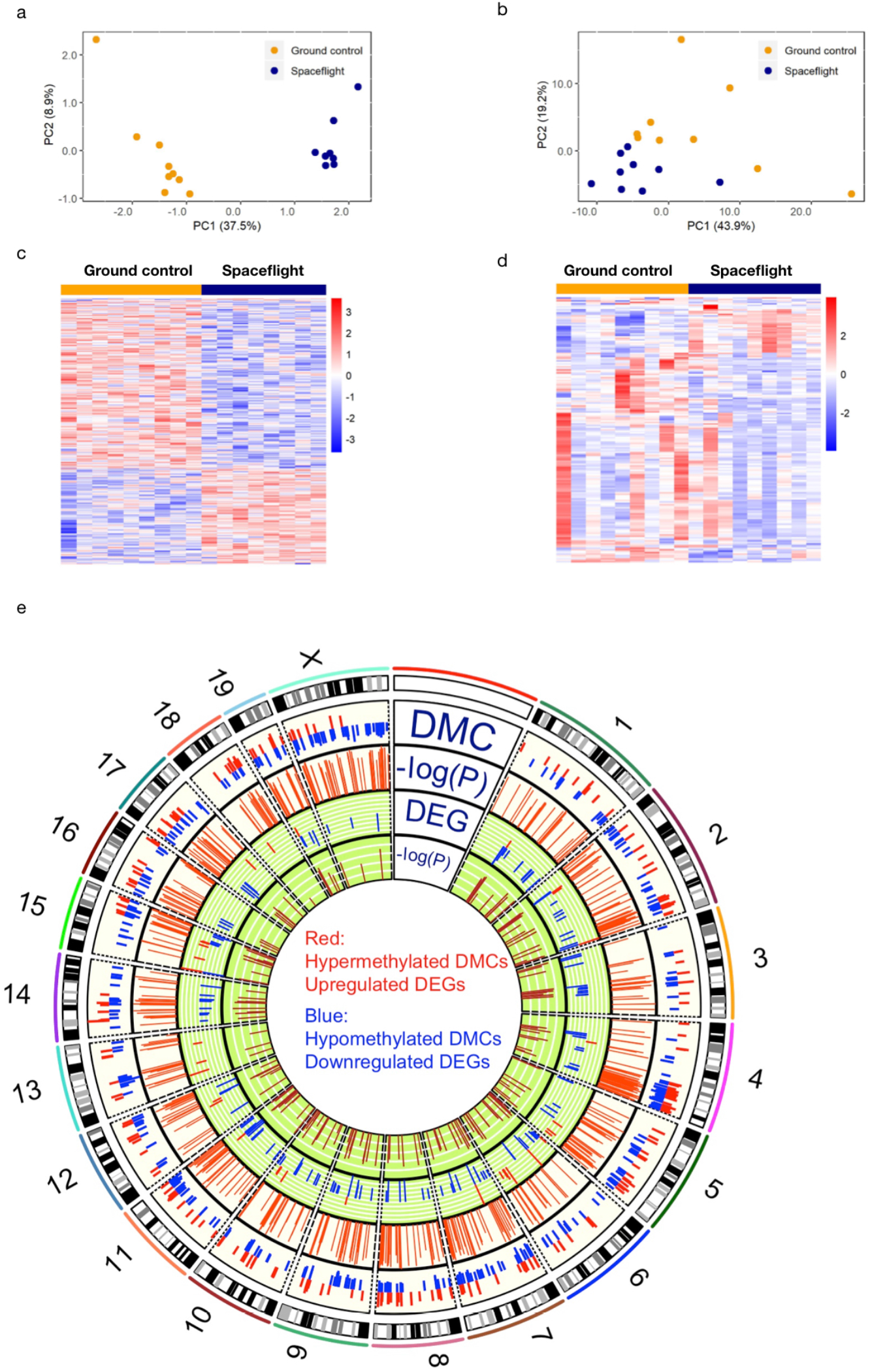
Spaceflight induced global alterations both on DNA methylation and gene expression. Principal component analysis (PCA) based on the spaceflight induced DMCs (**a**) and DEGs (**b**) in mouse retinas; Heatmap plots based on DMCs (**c**) and DEGs (**d**) between spaceflight mouse retinas and ground controls; (**e**) Circos plot showing the distribution of DMCs and DEGs across chromosomes. Y chromosome was excluded as only female mice were used for study.

Our RNA-seq transcriptomic analysis identified 177 DEGs (fold change ≥ 1.2 and p ≤ 0.05), including both up- and down-regulated genes (**Figure 2e and Supplementary Table 1**). Similar to that observed in methylome dataset, a distinct separation was observed between the spaceflight and ground control mice based on both the PCA and heat map of the DEGs (**Figure 3b and 3d**).

As DNA methylation has been successfully used to predict epigenetic clock—biological age^20^, we determined the epigenetic clocks of the retinas of spaceflight and control mice, using Stubbs’ epigenetic clock predictor^18^. The differences of Stubbs’ predicted biological ages between the spaceflight and ground control mice were tested using Welch two-sample t-test, which showed a significant age deceleration in spaceflight mice relative to ground mice (1.7 vs. 2.4 weeks, p = 0.049 on two-tailed t-test) (**Figure 2b**). It is worth noting that the predicted epigenetic ages were much younger than the true biological ages (21.2 weeks).

We further examined the alteration patterns due to the spaceflight on the DNA methylome and transcriptome across chromosomes using Circos plotting (**Figure 3e**). Despite that DMR was observed on each chromosome (chromosome Y was excluded as only female mice were used in this study), the DMRs were not evenly distributed across chromosome as well as at certain particular chromosome regions, suggesting that some chromosomes or regions were more prone to methylation modification. For instance, Chr X harbored the most DMRs, followed by Chr 4 and 7. In addition, some regions, e.g., Chr 4: 95278373-155343067 and Chr 2: 118262949-159068995, exhibited locally enriched DMRs. Although many DMRs were observed in certain chromosomes, e.g., Chr 2, 4, 5, 14, and X, there were fewer DEGs in these chromosomes (61 DMRs vs. 3 DEGs observed on Chr X; 30 DMRs vs. 14 DEGs observed on Chr 11), suggesting a less sensitive responsive in the gene expression of retinal tissue to spaceflight.

### 3. Biological functions and pathways affected by spaceflight in mouse retinas at transcriptome and DNA methylome, respectively

#### Functions and pathways affected in mouse retina by spaceflight based on DEGs

The IPA based on DEGs identified many significantly enriched canonical pathways (negative log [p-value] >1.3, i.e. P ≤ 0.05) due to spaceflight (**Figure 4a and Supplementary Table 2**). A large number of enriched pathways had a significantly negative Z-score (suppressed) and were associated with down-regulated DEGs (**Figure 4a**). Enriched canonical pathways mainly belonged to two biological processes: 1) the junction/connection including G6P signaling, osteoarthritis signaling, and ILK signaling pathways, and 2) the inflammation/proliferation including PDGF signaling, CXCR4 signaling, actin cytoskeleton signaling, and NF-κB signaling. Many other biological pathways, such as hepatic fibrosis, IGF-1 signaling, epithelia adheren junction, and tight junction (**Supplementary Table 2**), were enriched by spaceflight with significant p values, but zero Z-scores due to a lack of other relevant supporting evidence. These pathways were functionally linked to the extracellular matrix (ECM) homeostasis. Notably, many of the above pathways enriched and impacted by spaceflight in this study were also observed in NASA Twins Study^16^. At individual gene level, a large number of DEGs fell into the categories of wound healing/cell growth/cell motility, inflammation defense, mitochondria metabolism, and oxidative stress (**Table 1, Figure 4a**). In summary, the IPA analysis indicated impaired cell-to-cell connection and subsequent repair.

**Figure 4.**
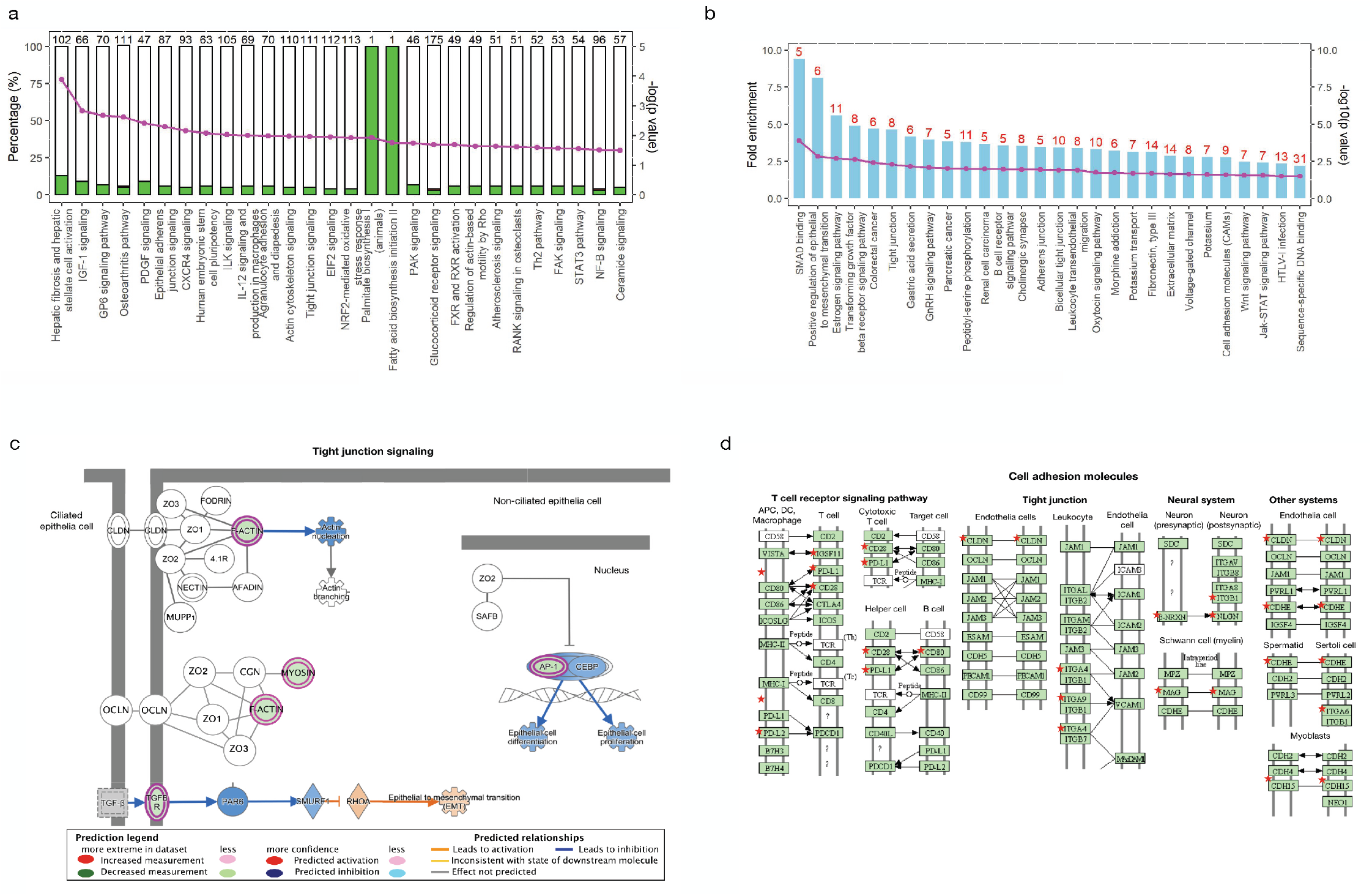
Functional enrichments based on the spaceflight induced DEGs and DMGs. (**a**) The IPA identified significantly enriched canonical pathways based on spaceflight induced retinal DEGs between spaceflight mice and ground controls. The left y-axis shows the percentage of DEGs to the total number of genes in each individual pathway; the white unfilled bar section represents the percentage of non-DEGs, the filled green bar section represents the percentage of down-regulated DEGs, and the red filled bar section represents the up-regulated DEGs. The right y-axis shows the negative log_10_ (p-value) of the enriched pathways as pink dot-line. The negative log_10_(0.05)=1.3. (**b**) Significant pathways enriched based on differentially methylated genes (DMGs) using the DAVID Functional Annotation Analysis. The bars represent the fold enrichment and the dot-line represents the negative log_10_(p-value). Red number on top of each bar indicates the number of DMGs identified in each pathway. (**c**) Tight junction signaling, a representative pathway enriched based DEGs. Pathway components are highlighted by blue (down-regulated) or purple (up-regulated) showing the DEGs identified within the component. (**d**) Cell Adhesion Molecules, representative functional clusters enriched by DAVID functional analysis. The significant genes identified in our spaceflight mouse retinal methylome were labelled as red star.

In addition, based on the functional similarity, the DAVID analysis identified 47 genes out of 177 DEGs as secreted/extracellular proteins, and 17 genes of those 47 DEGs were the EMC components (**Table 2**). Furthermore, transforming growth factor beta (*Tgfb*) signaling, wound healing process, and insulin-like growth (IGF) factor signaling were enriched by 12, 9.9, and 6 folds, respectively, in spaceflight mouse retinas as compared to those housed in AEM (**Table 2 and Supplementary Table 1**, P≤0.01). Several genes enriched in the TGF-β group, such as *Pdgfra, Nog,* and *Dcn*, were significantly down-regulated (P≤0.05, N = 9 per group). The transcriptional factors such as hormone-retinoid receptors *Nr4a1, Nr4a3*, zinc finger transcriptional factor *Egr1*, and connective tissue growth factor Ctgf were observed to be down-regulated in spaceflight mouse retinas (**Table 1 and Supplementary Table 1**). In contrast, the majority of the top up-regulated DEGs were the structural protein genes, like *Krt1*, *Krt10*, and *Adamts3* gene, which is consistent with the notion that spaceflight activated inflammation and cell survival mechanisms. In summary, consistent with the IPA analysis, the David functional analysis suggested that spaceflight compromised cell proliferation and mobility necessary for retinal wound healing.

#### Functions and pathways affected in mouse retina by spaceflight based on DMGs

We identified 434 hypomethylated and 209 hypermethylated genes, using 10% difference on methylation and P<0.05 (n=9 for each group) as criteria (**Figure 2c and Supplementary Table 1**). Out of those 643 DMGs, 118 had at least one differentially methylated cytosine (DMC) located within a CpG island and 108 had DMC located in the promotor regions (34 hypomethylation and 74 hypermethylation) (**Figure 2d**), indicating epigenetic regulation of transcriptional changes in spaceflight compared to ground control mice. Furthermore, 22 hypermethylated genes and 35 hypomethylated genes had DMCs located in both CpG islands and exon areas.

We used the DAVID Functional Clustering to examine how spaceflight induced methylome changes would affect retinal cell function. The data revealed that a substantial amount of DMGs were link to cell structure and tissue morphology, with significant p values and fold enrichment (**Figure 4b and Supplementary Table 2**). Specifically, 35 DMGs were related to the cell junction, 17 DMGs were related to the extracellular matrix, and 8 DMGs were related to the tight junction. Another 20 DMGs were members of collagen and fibronectin gene families, with fold enrichment values of 3.4 (P<0.05) and 3.1 (P<0.01), respectively. Further, functional clustering revealed that DNA methylation might play an important role in cell proliferation, differentiation, and organism development (**Table 2 and Supplementary Table 2**). For instances, 24 DMGs were linked to cell differentiation and 38 DMGs were linked to multicellular organism development. In addition, 6 DMGs (fold enrichment value of 8.1, P<0.01) were positive regulators of epithelial to mesenchymal transition and several other DMGs were components of Hippo and TGF-β signaling pathways. DMGs functional analysis demonstrated consistent shifts in cell function and agreed well with those observed in transcriptome analysis.

DMGs based IPA analysis portrayed a concerted hypermethylation on several retinoic acid (RA) signaling pathway genes **(Supplementary Figure 2)**. Among those were: *Rho* (rhodopsin; light perception), *Vegfa* (vascular endothelial growth factor A), *Tgfb1* (cell growth factor), *Bcl2l11* (apoptosis facilitator), *Mmp2* (matrix metallopeptidase 14), *Wnt5a* (transcription factor), and *N32e1* (retinal transcription factor). Consistent with the changes in RA signaling, *Nxph1* gene, which is involved in neuronal synaptic transmission, was highly methylated. Genes with decreased methylation included: *Smad2*, *Smad3*, *Sod3*, *Tnfrsf19*, *Edn2*, and *Dnmt3l* **(Supplementary Table 2)**. Six mitochondrial membrane solute transporter genes (*Slc22a2, Slc25a25*, *Slc25a33, Slc35f3, Slc6a17*, and *Slc6a8*) were hypomethylated, suggesting that mitochondrial function was affected during spaceflight.

Further functional analysis suggested that spaceflight could induce morphological changes in many retinal cell types. The putative morphology changes were evidenced by the extensive functional overlaps of 9 DMGs (5 hypermethylated: *Erc2, Neurog2, ATP8A2, App, and Rarg; 4 hypomethylated: Rho, Vegfa, Hes1, and Discam*) **(Supplementary Table 2)**. Particularly, enrichment analysis implicated that at least two DMGs were involved in the abnormal morphological development in horizontal cells, photoreceptor cells, retinal layer cells, and retinal rods on the photo-receptor side and bipolar cells and neurons on the transmitter side.

### 4. Integration analysis of genes regulated at both transcription and DNA methylation levels

We identified 5 genes in the mouse retinas regulated by spaceflight that were present both in the DEG and DMG lists: *Col8a2, Jun/Juneb, Maf, Mid1ip1*, and *Tsc22d3* (**Figure 5a-b, and Supplementary Table 1**). It is worth noting that *Mid1ip1*, a cellular growth regulator, was hypomethylated in its promoter, exon, and CpG island regions, but its transcription was down-regulated (**Figure 5b**). Interestingly, although *Col8a2* (a collagen gene), *Maf* (a transcription factor), and *Junb* (a transcription factor) had significant changes in both transcription and DNA methylation, the differentially methylated CpGs were far from their gene bodies (**Supplementary Table 1**). Hypomethylation was found in the intron and CpG island regions for *Tsc22d3*, whereas its transcription was suppressed (**Figure 5b**).

**Figure 5.**
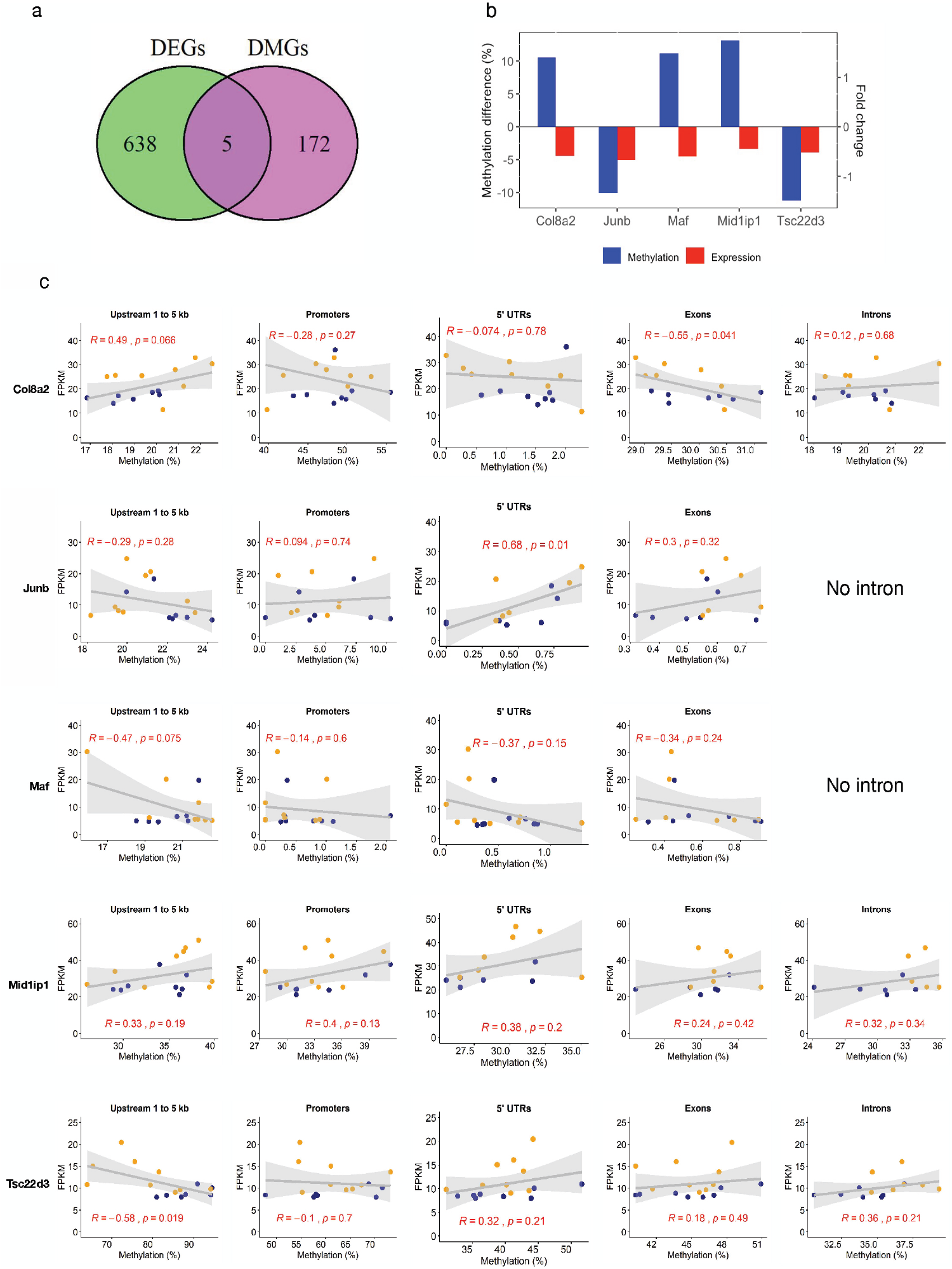
Integrative analysis of spaceflight induced DMGs and DEGs. (**a**) Venn diagram showing the overlapping DMGs and DEGs induced by spaceflight, (**b**) The DNA methylation (based on differentially methylated CpG sites of the gene) vs. the gene expression for the five common DMGs and DEGs. The left y-axis shows the DNA methylation change (blue bar for DNA methylation); the right y-axis shows the gene expression fold-change (red bar for gene expression). (**c**) Correlation between the gene expression and DNA methylation of different location of gene for the five overlapping genes. Blue, spaceflight mouse retina; orange, ground control mouse retina.

In order to understand the regulatory role of DNA methylation on transcription, we performed Pearson correlation analysis between DNA methylation and gene expression at gene body level, using all CpGs annotated to the 5 overlapping genes (**Figure 5c**). Overall, there were mild to moderate correlations (|R| values were between 0.3 ~ 0.7), nevertheless with no consistent trend between DNA methylation in the gene body and their gene expressions. For examples, in Col8a2 gene, a moderate positive correlation between DNA methylations at upstream 1 to 5 kb region (R=0.49, p=0.066) but a moderate negative correlation (R=−0.55, p=0.0410) at exons and its gene expression were observed. In *Junb* gene, a moderate positive correlation (R=0.68, p=0.01) between DNA methylation at 5’ UTR and its gene expression was observed. In *Tsc22d3* gene, a moderate negative correlation (R=−0.58, p=0.019) was observed between DNA methylation at upstream 1 to 5kb region and its gene expression. Weak negative correlations were observed between DNA methylations in all gene body regions of *Maf* and its gene expression, while weak positive correlations were observed between DNA methylations in all parts of *Mid1ip1* gene and its gene expression.

### 5. Genes and pathways regulated by spaceflight at both transcription and DNA methylation levels

The functional analysis based on both DEGs and DMGs revealed that the genes involved in ECM and focal adhesion were the main regulatory targets in the retinas by spaceflight. We found that ECM genes like collagen, laminin, thrombospondin (*Thb*s), and integrins included *CD44*, syndecan, and VLA protein β subunit 1, 3, 8, were regulated at transcriptional level, whereas other genes like tenascin and integrin VLA α subunit 2, 7 and 8, were regulated at DNA methylation level by spaceflight (**Supplementary Figure 3a**). Though simultaneously changes at both the transcription and DNA methylation levels were not observed in many genes, an integration analysis identified many DEGs and DMRs were enriched in the ECM/cell junction functional group, suggesting that the genes and gene networks in the ECM/cell junction were more sensitive to the stress of spaceflight in the mouse retinas. This notion of convergent regulation is illustrated in **Figure 4c** and **4d**.

We examined the DEGs and DMGs involved in HIPPO, TGF-β, and Wnt signaling pathways (**Supplementary Figure 3b**). Our analysis showed that *Fzd*, a transmembrane receptor in Wnt pathway, was significantly downregulated in gene expression; while *Tfg-βr*, a transmembrane receptor in TGF-β pathway, was hypermethylated. *Fzd* and *Tfg-βr* interact with a variety of extracellular signal molecules to activate signaling cascade, of which many downstream signal components were found in either the DEG or DMG lists, including the genes responsible for apoptosis and proliferation*, such as Smad2/3, CK1δ/∊, Axin, CycD,* and *αPkc*. Furthermore, the epithelial adheren, a cell surface receptor for calcium-dependent cell-cell adhesion, was hypomethylated. Overall, the integrated gene network and pathway analysis strongly suggests a coordinated response at the transcriptome and methylome levels in the spaceflight mice.

## DISCUSSION

Approximately one in three astronauts flying on long-duration space missions experience visual impairment and morphologic changes to their eyes that include choroidal and retinal folds, optic disc edema, focal areas of retinal ischemia (i.e., cotton wool spots), globe flattening, and hyperopic shifts. This collection of ocular disorders, as first reported by Mader and colleagues^3^, has been termed spaceflight-associated neuro-ocular syndrome (SANS). A number of factors associated with spaceflight have been proposed to initiate SANS, including headward fluid shifts induced by weightlessness, space radiation, and high ambient CO_2_ levels within the spacecraft. A variety of potential mechanisms have also been proposed to account for the unusual physiologic and pathologic neuro-ophthalmic findings, including elevations in intracranial pressure (ICP) from cephalad fluid shifts, altered autoregulation of cerebral perfusion, impaired cerebrospinal fluid drainage from the brain and orbital optic nerve sheath through venous, glymphatic, and lymphatic drainage systems, and disruption of blood-brain, blood-retinal, and blood-optic nerve barrier function^21^. This seemingly multifaceted pathological process, which varies from astronaut to astronaut, indicates a complex origin for these neuro-ophthalmic findings associated with SANS. Results from the present study of spaceflight-induced alterations in DNA methylome and transcriptome demonstrate that retinal cell homeostasis is disrupted during spaceflight, and that the primary impacted genes belong to several physiologically relevant cellular processes and pathways. These include processes and pathways associated with oxidative stress, inflammation, mitochondrial function, tissue remodeling, fibrosis and angiogenesis. Further, the integrated DNA methylome and RNA transcriptome analysis demonstrates that spaceflight had profound effects on ECM/cell junction and cell proliferation/apoptosis signaling in the retina. Although these data do not address all the possible mechanisms involved in the etiology of SANS, they provide crucial insight into the potential adverse consequences of spaceflight on the retina that could be functionally important for maintaining proper visual acuity among astronauts.

Multi-dimensional genome-wide analyses have been applied to interrogate vision function, aging and diseases^22^. In this study, we characterized the retinal tissue epigenomic reprogramming, along with the transcriptomic changes of mouse retinas after a period of 37-day spaceflight, the human equivalent of 4.1 years. The differential gene expression analysis revealed a few distinctive yet interconnected pathways that may have been affected by spaceflight, such as inflammatory and cancer-prone pathways, i.e., interleukin and chemokine pathways (*Fgfr1, Fos, Cxcr4* genes); cell structure maintenance pathways, i.e., ECM and keratins; and the cell death, cell differentiation, and survival pathways, i.e., VEGF, TSP1, TGF-β signaling and DNA repair. These findings support the notion that environmental conditions during spaceflight, such as microgravity and space radiation, induce retinal structure damage and the counteractive cell response^8^, leading to similar conditions of known disease states such as age-related macular degeneration (AMD)^15,23^.

Consistent with what has been reported on retinal disorders^13^, we identified six DEGs in the spaceflight mouse retinas that were associated with AMD (*Adamts, Col8a, Htr, Igfb, Slc, Tgfbr*). We also found that the mitochondrial small molecule transporter family genes such as *Slc4a7, Slc6a17, Slc22a8, Slc22as, Slc25a2,* and *Slc31a2* were among the top DEGs and DMGs induced by spaceflight in the mouse retinas. A gene expression change of a small molecule transporter in ocular tissue was also observed in a previous spaceflight study, which was linked to oxidative damage in leading to mitochondrial apoptosis and cell death in retina^8^. Many other structural proteins, such as collagen, keratin and actin, were also downregulated under spaceflight conditions. The suppression of these genes has been shown to trigger chronic inflammation, fibrosis, retinopathy, and cancer^24^. Notably, several genes in the retinoic acid metabolism pathway were hypermethylated in spaceflight mice. RA receptors repress the activity of transcription factors such as *AP-1* and *NF-κB*, and therefore have potential anti-proliferative and anti-inflammatory effects. In addition, the expressions of several genes (*Wnt5a, Vegf, and Tnf*) in Wnt/β-catenin pathway were altered by spaceflight. Wnt/β-catenin pathway plays a pathogenic role in AMD, and it is believed that certain changes in AMD likely lead to inflammation or a cancer-like disease state^25^.

Intraocular pressure (IOP) impacts cell adhesion and connection, promotes detachment of cell from ECM, leading to apoptosis and optical nerve damage^26,27^. Our transcriptomic and epigenomic studies on spaceflight mice retinas showed that ECM-receptor components were among the most altered genes in spaceflight mice retinas relative to that in ground control mice (**Table 1**). We found that the expression of myocilin (encoded by *Myoc*), a structural component that regulates IOP, was significantly suppressed, suggesting an adverse effect of microgravity on IOP regulation. Another significantly altered gene family in the spaceflight mouse retinas was simple epithelial keratins (SEKs), which are primarily expressed in single-layered simple epithelia. An important feature of SEKs is that one specific SEK pair (one type I SEK and one type II SEK) predominate in an epithelial cell where they form noncovalent heteropolymers^28^. Our transcriptomic data showed both *Krt1* and *Krt10* were significantly upregulated, suggesting that the spaceflight mice might have been subject to some structural repair in the retinas. However, the expression of another corneal epithelial SEK*, Krt12*, was significantly suppressed. Beside its structural roles, *Krt12* is generally regarded as a specific corneal epithelial differentiation marker^29^ and a reduced expression of *Krt12* gene may suggest an impaired cell differentiation under spaceflight condition.

The transcriptomic data revealed spaceflight down-regulated wound-healing and cell proliferation genes (*Tgfb2, Tgfb3, Igfbp4, Igfbp5, Igfbp6*, *Pdgfra, Dcn*) in mouse retinas. Furthermore, many transcriptional regulators were also downregulated in the spaceflight mouse retinas, including hormone-retinoid receptors *Nr4a1* and *Nr4a3*, zinc finger transcriptional factor *Egr1*, connective tissue growth factor *Ctgf*, and serine protease inhibitors *Serpine2, Serpinf1, Serpinh1* and *Serping*. These transcriptional factors play critical roles in regulating cell survival, proliferation, axial eye growth^30^, morphogenesis of the lens and optic vesicle development^31,32^. The methylomic data also suggests that spaceflight may impact cell cycle and DNA repair as indicated by the IPA analysis based DMGs, which showed that R-SMAD binding protein, TGFBR signaling, and WNT signaling pathways were all enriched. The recent NASA Twins Study^16^ also showed that the gene expression changed for genes related to cell growth, proliferation and angiogenesis in CD4 and CD8 cells, indicating that the activation of wound-repairing pathways are not limited to retinal tissue in spaceflight.

This is the first report on an epigenetic age alteration in retinal tissue during spaceflight. The data indicated that the retinas of spaceflight mice tended to have a younger epigenetic age (deceleration of age) compared to that in ground control animals (**Fig. 2b**). Interestingly, in the NASA Twin Study, the results showed that the spaceflight twin had an elongated telomere length relative to his ground-based twin brother, indicating slower aging in space^16^. Thus, there seems to be some evidence that spaceflight-induced age deceleration occurs in mice and humans. There also seems to be some similarity in the aging deceleration of the retinal epigenetic clocks of spaceflight mice and that in mice with ovariectomy or on a high-fat diet^18,19^. It is worth noting that the predicted epigenetic age based on retinal methylation was much younger than the chronological age (2~3 weeks as predicted by retinal DNA methylation clock vs. 21 weeks per chronological age). A couple of factors might have contributed to this discrepancy. First, the retinal tissue DNA methylation data was not included in Stubbs’s epigenetic clock training dataset^18^, and therefore, the retinal epigenetic age was not well represented by Stubb’s algorithm. Second, the retina is a heterocellular tissue and its epigenetic age might be over-represented by epigenetically younger cells. As matter of fact, Horvath reported that several tumors exhibited decelerated epigenetic clocks and that malignant cancer tissues were much younger than anticipated^17,33^. Nevertheless, our spaceflight mouse retinal epigenetic clock analysis showed that spaceflight significantly decelerated retinal epigenetic age. This age deceleration may suggest an activation of remodeling or compensatory pathways that are dormant under normal physiological conditions^34^.

In summary, this study is the first to investigate the effects of spaceflight on both DNA methylome and transcriptome in retinal tissue. Interestingly, the results indicate that spaceflight decelerated epigenetic clocks in the mouse retina, i.e., the mice flown in space showed younger biological ages than the ground control mice. Despite this positive finding, the results also demonstrated that retinal cell homeostasis was disrupted during spaceflight. The primary impacted genes by spaceflight belong to several physiologically relevant cellular processes and pathways, including those associated with oxidative stress, inflammation, mitochondrial function, tissue remodeling, fibrosis and angiogenesis. The functional analysis integrating both DNA methylome and RNA transcriptome demonstrated that spaceflight had particularly profound effects on two important biological processes, i.e., ECM/cell junction and cell proliferation/apoptosis signaling in the retina (**Figure 4c, 4d**, **and Supplementary Figure 3**).

## Methods

### 1. Study design

**Figure 1**. illustrates the overall study design. The 16-week-old female C57 BL/6J mice (Jackson Lab, Bar Harbor, ME) were used in this study as part of the NASA RR-1 project consortium; this project was approved by the NASA Institutional Animal Care and Use Committee. Both spaceflight and ground control animals were housed in NASA’s animal enclosure modules (AEM), with control mice being exposed to the same environment conditions (12-hour light cycle, temperature and humidity) as those flown on the International Space Station (ISS). Control animals were kept inside an environmental simulator (ISSES) at the Space Life Science Laboratory (SLSL) at Kennedy Space Center (KSC), and the spaceflight animals were transported to the ISS by SpaceX4 on September 21, 2014. All animals were fed with a special NASA food bar diet and their health were checked daily. The spaceflight mice were euthanized and frozen on orbit after 37 days of flight, while ground control mice were simultaneously euthanized and frozen under identical conditions. After the frozen carcasses were returned to KSC, the ocular tissues were removed from both groups. DNA and RNA were extracted from the same retinal sample and were used for library constructions using NuGEN Ovation^®^ RRBS and Ovation^®^ Mouse RNA-seq library construction kits, respectively. Libraries were multiplexed with unique sample indices, pooled, and sequenced on an Illumina NextSeq 550. FastQ files were subject to FastQC, adapter and low-quality reads trimming before mapping to mouse genome NCBI GRCm38. Differential CpG methylation regions (DMRs) and differential expressed genes were determined after mapping and counting, and the resulting DMR and DEG lists were subject to IPA and DAVID gene functional analysis. The epigenetic clock was determined using the MouseEpigeneticClock script developed by Stubbs et al. (2017)^18^.

### 2. Dissection and preservation of the mouse ocular samples

At the time of dissections, both eyes were dissected from the carcasses thawed at room temperature for 15-20 minutes. The eyes were harvested and snap frozen in liquid nitrogen, then stored at −80 °C. The frozen whole eye samples were micro-dissected in the ice-cold phosphate-buffered saline (PBS). Fine spring scissors were used to separate the cornea from the sclera to access the retina. The retina was carefully detached from the sclera. The retina, cornea, and lens were immediately frozen on dry ice and stored at −80°C before DNA and RNA isolations.

### 3. DNA and RNA isolations

Retinal tissues (4-20 mg) were homogenized with Precellys CKM beads (Hilden, Germany) in 350 μl of Qiagen RLT plus buffer. The genomic DNA (gDNA) and total RNA were isolated using the Qiagen AllPrep DNA/RNA/miRNA Universal Kit according to the manufacturer’s protocol. After isolation, RNA and DNA were frozen and kept at −80°C until further use. Isolated RNA samples were quantified using NanoDrop spectrophotometry (Thermo Scientific, Chino, CA). DNA and RNA were further quantified using the Qubit dsDNA High Sensitivity Kit and RNA Broad Range Kit, respectively (Life Technologies, Carlsbad, CA). RNA quality was evaluated using the Agilent 2200 TapeStation and RNA ScreenTape (Santa Clara, CA). All RNA samples had an integrity numbers (RIN) ranging from 7.2 to 8.1.

### 4. Reduced Representation Bisulfite Sequencing (RRBS) library construction

One hundred (100) ng of gDNA was used to construct RRBS DNA-seq library using the Ovation^®^ RRBS Methyl-Seq System (NuGEN Technologies, San Carlos, CA) according to the manufacturer’s protocol. Briefly, the MspI enzyme, which cuts the DNA at CCGG sites, was used to digest gDNA into fragments. The fragments were directly subject to end blunting and phosphorylation in preparation for ligation to a methylated adapter with a single-base T overhang. A unique index, out of 16 indices, was used for each individual sample for multiplexing. The ligation products were final repaired in a thermal cycler under the program (60°C – 10 min, 70°C – 10 min, hold at 4°C). The products of the final repair reaction were used for bisulfite conversion using kit QIAGEN EpiTect Fast DNA Bisulfite Kit according to Qiagen’s protocol. Bisulfite-converted DNA was then amplified (Mastercycler^®^ pro, Eppendorf, Hamburg, Germany), and bead-purified with Agencourt RNAClean XP beads.

### 5. RNA-seq library construction

The Ovation^®^ Mouse RNA-Seq System 1-16 (NuGEN Technologies, San Carlos, CA) was used to construct RNA-seq libraries according to the manufacturer’s instructions. Briefly, 100 ng of total RNA spiked with 1 μl of 1:500 diluted ERCC Mix 1 (Life Technologies, Carlsbad, CA) was used to start cDNA synthesis. Following the primer annealing and cDNA synthesis, the products (130 μl/each sample) were sheared using Covaris S220 Focused-ultrasonicator (Covaris Inc., Woburn, MA). The parameters were set as follows: 10% duty factor, peak power 175 and 200 cycles per burst at 4°C for 200 seconds to obtain fragment sizes between 150-200 bp. The products were then subject to end-repair, adaptor index ligation and strand selection. A custom InDA-C primer mixture SS5 Version 5 for mice was used to allow strand selection. Finally, libraries were amplified on a PCR thermocycler for 17 cycles (Mastercycler^®^ pro, Eppendorf, Hamburg, Germany), and purified with RNAClean XP Agencourt beads (Beckman Coulter, Indianapolis, IN); and quantified using Qubit dsDNA HS Kit on Qubit 3.0 Fluorometer (Life Technologies, Carlsbad, CA). The quality and peak size were determined using the D1000 ScreenTape on Agilent 2200 TapeStation (Agilent Technologies, Santa Clara, CA).

### 6. RRBS and RNA-seq library sequencing

The RRBS libraries were sequenced on an Illumina NextSeq 550, 75 bp, single-end read, whereas the RNA-seq libraries were sequenced on both Illumina NextSeq 550 and HiSeq 4000, pair-end reads (75 bpx2), at the Loma Linda University (LLU) Center for Genomics.

### 7. Bioinformatics pipelines

For the RRBS data, we used a pipeline that integrates the read quality assessment (FastQC), trimming process (TrimGalore^35^, NuGEN diversity trimming and N6 de-duplicate scripts), alignment (Bismark^36^), and differential methylation analysis using MethylKit^37^ and DMAP^38^. This pipeline facilitates a rapid transition from sequencing reads to a fully annotated CpG methylation report for biological interpretation. Briefly, the RRBS raw fastq data were first trimmed using Trim Galore. The mouse genome NCBI GRCm38 downloaded from iGenome (https://support.illumina.com/sequencing/sequencing_software/igenome.html/Musmusculu/Mus_musculus_NCBI_GRCm38.tar.gz), was used as a reference genome. Reads were aligned to the mouse reference genome with Bismark v0.16.334 by default parameter settings. The methylation call files including the location of each CpG site and the methylation percentage were generated by the bismark_methylation_extractor function. The aligned SAM files were further processed through DMAP^38^ to generate CpG region profiling. The CpG regions with coverage by a minimum 20 reads in all samples were used for follow-up analysis.

For the RNA-seq data, we adopted the pipelines used in our recent publications^39,40^ for mRNA-seq data visualization, which integrated the QC (FastQC^41^, ShortRead^42^), trimming process (trimmomatic^43^), alignment (Tophat2^44^), reads quantification (cufflinks^45^) and differentially expressed gene (DEG) analysis (cuffdiff^46^) for mRNA-seq data analyses. Briefly, the RNA-seq raw fastq data were first trimmed using Trimmomatic. The trimmed reads were aligned to the mouse reference genome (NCBI GRCm38) using TopHat V2.1.1 with default parameter settings. The aligned bam files were then processed using Cufflinks V2.2.1 for gene quantification. The reads that were unable to align to the mouse genome were converted to fastq format using SamToFastq function in Picard V1.114 for ERCC mapping and calculation in which the reads were mapped to ERCC transcripts, and quantified using TopHat V2.1.1 and Cufflinks V2.1.1 with default parameter settings. The genes with FPKM ≥ 1 in all samples were used for DEG analysis. The differentially expressed genes (DEGs) were identified by Cuffdiff with FDR ≤ 0.3, and fold change (FC) ≥ 1.2 (P ≤ 0.05).

### 7. Epigenetic clock calculation

We used a mouse epigenetic clock algorithm developed by Stubbs et al. (2017)^18^ to estimate the biological age of the retinas of spaceflight mice, using the MouseEpigeneticClock script on the methylation coverage files with default parameter of read depth threshold at 5. The mouse age estimator, based on the methylation of 329 CpG loci, was shown to accurately measure the epigenetic clock—biological age across multiple mouse tissues^18^.

### 8. Gene networks, pathway and functional annotation analyses

The analyses on the gene networks, canonical and bio-functional pathways regulated by spaceflight at both DNA methylome and transcriptome were performed using the Ingenuity Pathway Analysis (IPA, Qiagen). The sets of 177 differentially expressed genes and 643 differentially methylated genes were input into the IPA database to discover canonical, diseases and bio-functional networks possibly regulated by spaceflight. The complete IPA analysis summary is shown in the supplementary files. We also applied the DAVID Functional Annotation Tool (https://david.ncifcrf.gov) to the lists of DEGs and DMGs to identify the functional enrichment of the spaceflight-induced gene expression changes to biologically relevant categories.

### 9. Statistical analysis

For DEGs analysis, a FC ≥ 1.2 with P ≤ 0.05 (FDR ≤ 0.3) was used as a threshold to select DEGs. For DMG analysis, a methylation change ≥10% (FDR ≤ 0.05) was used as a threshold to select DMGs. Pathway analysis was performed using IPA and DAVID Functional Annotation Tool to identify canonical pathways. For other statistics analysis, data were presented as mean ± SD, and were analyzed using two-tailed Student’s t-test or one-way ANOVA followed by Turkey’s post hoc test. P-value ≤ 0.05 was considered statistically significant. Experimental number (N) represents number of animals tested in each group.

## Data availability

The RRBS and RNA-seq sequencing data has been uploaded to the Gene Expression Omnibus (GEO) with the accession GSE159771. The data remain private currently and can be accessed when the paper is published. The following is the reviewer’s link and a private access token can be obtained from the journal editor: https://www.ncbi.nlm.nih.gov/geo/query/acc.cgi?acc=GSE159771

## Author contributions

CW, XWM and MDD conceived the study and CW designed and determined the experimental approach. MDD and XWM obtained the eye tissue through the NASA Space Biology Program. ZC, SS carried out the experiments, performed bioinformatics data analysis and drafted the manuscript. NCN performed the mouse retina isolation. XC performed bioinformatics data analysis. XC, MDD, HQ and XWM reviewed the manuscript. CW revised and finalized the manuscript.

## Acknowledgements

The authors would like to thank Dr. Thomas M. Stubbs, Dr. Wolf Reik and Dr. Zhaowei Yang for the discussion and help in the analysis with the mouse epigenetic age estimator. The genomic work carried out at the Loma Linda University Center for Genomics was funded in part by the National Institutes of Health (NIH) grant S10OD019960 (CW). This project was partially supported by NASA Space Biology grant NNX15AE86G (MDD and XWM) and AHA grant 18IPA34170301 (CW); and also partially supported by NIH grants HL115195-06 (HQ)/subcontract (GSU) # SP00013920-02 (CW), and HL137962 (HQ)/subcontract (GSU) # SP00013696-01 (CW).

