## Supplementary Figures for "Spaceflight decelerates the epigenetic clock orchestrated with a global alteration in DNA methylome and transcriptome in the mouse retina"

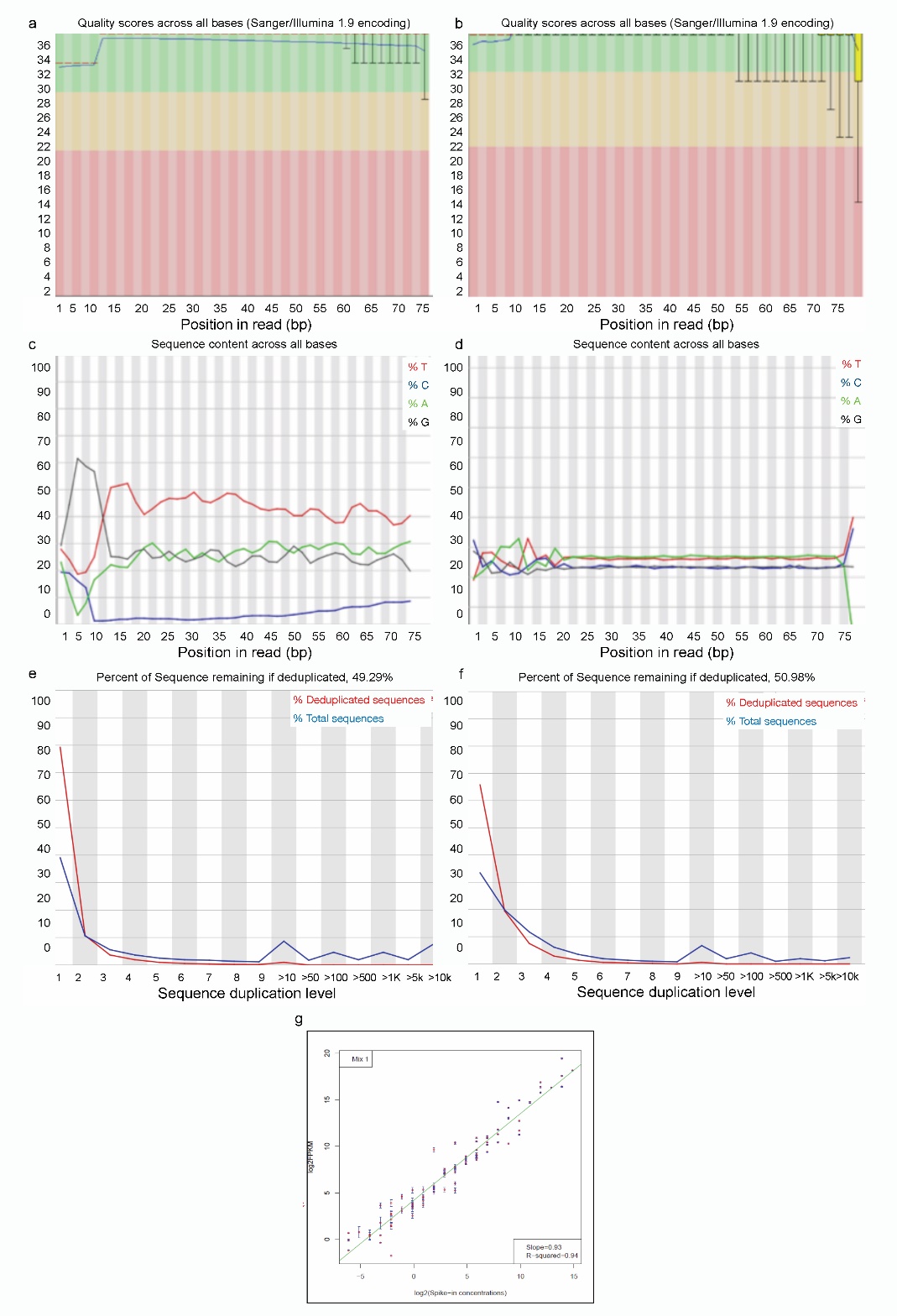

**Supplementary Figure 1: Sequencing data quality control.** Fastq file quality of a representative RRBS read (**a, c, e**) and an RNA-seq (**b, d, f**) sequencing read across bases. (**a**, **b**) Box and Whisker plots showing the Phred quality scores at each position of bases. The central red line is the median value. The yellow box represents the inter-quartile range (25–75%). The upper and lower whiskers represent the 10% and 90% points. The blue line represents the mean quality. (**c**, **d**) Per base sequence content after adaptor sequence removal, (**e**, **f**) Sequence duplication level. The plot shows the proportion of the library which contained sequences in different duplication level bins. The blue line shows the total sequences and the duplication level distribution. The red line shows the proportions of sequences after e-duplication. (**g**) Scatterplot showing the correlation of RNA-seq between the detected ERCC log2 (FPKM) and the expected ERCC log2 (spike-in concentration).

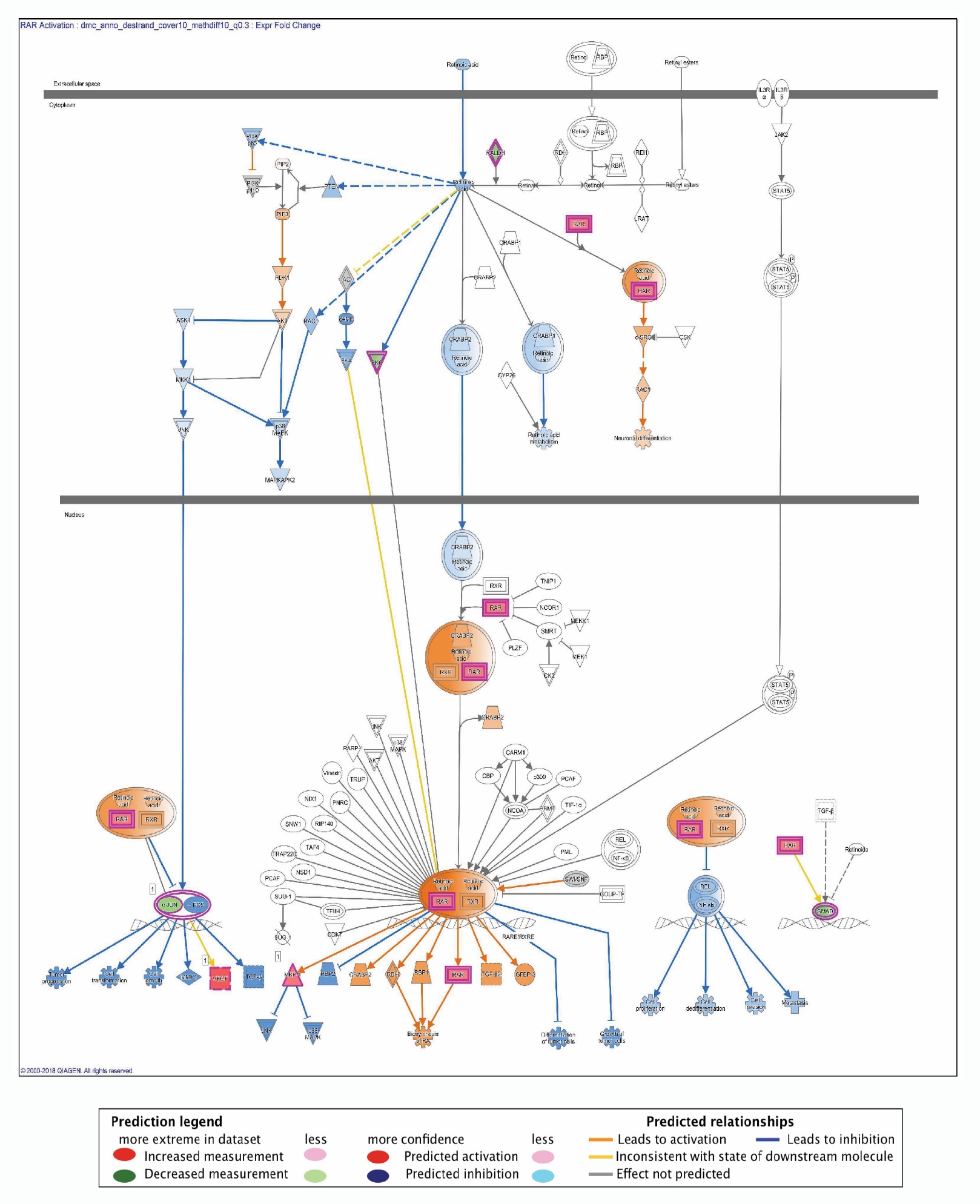

**Supplementary Figure 2.** **DMGs identified in the retinoic acid receptor (RAR) pathways**. Molecules identified in this pathway were Jun, Prkcd, Dusp1, Vegfa, Smad2, Smad3, Rarg, Aldh1a3, and Prkcz. Prediction of downstream and upstream molecules relations from the identified molecules are depicted (refer to the Prediction Legend). Canonical pathways generated by Ingenuity Pathway Analysis based DMGs.

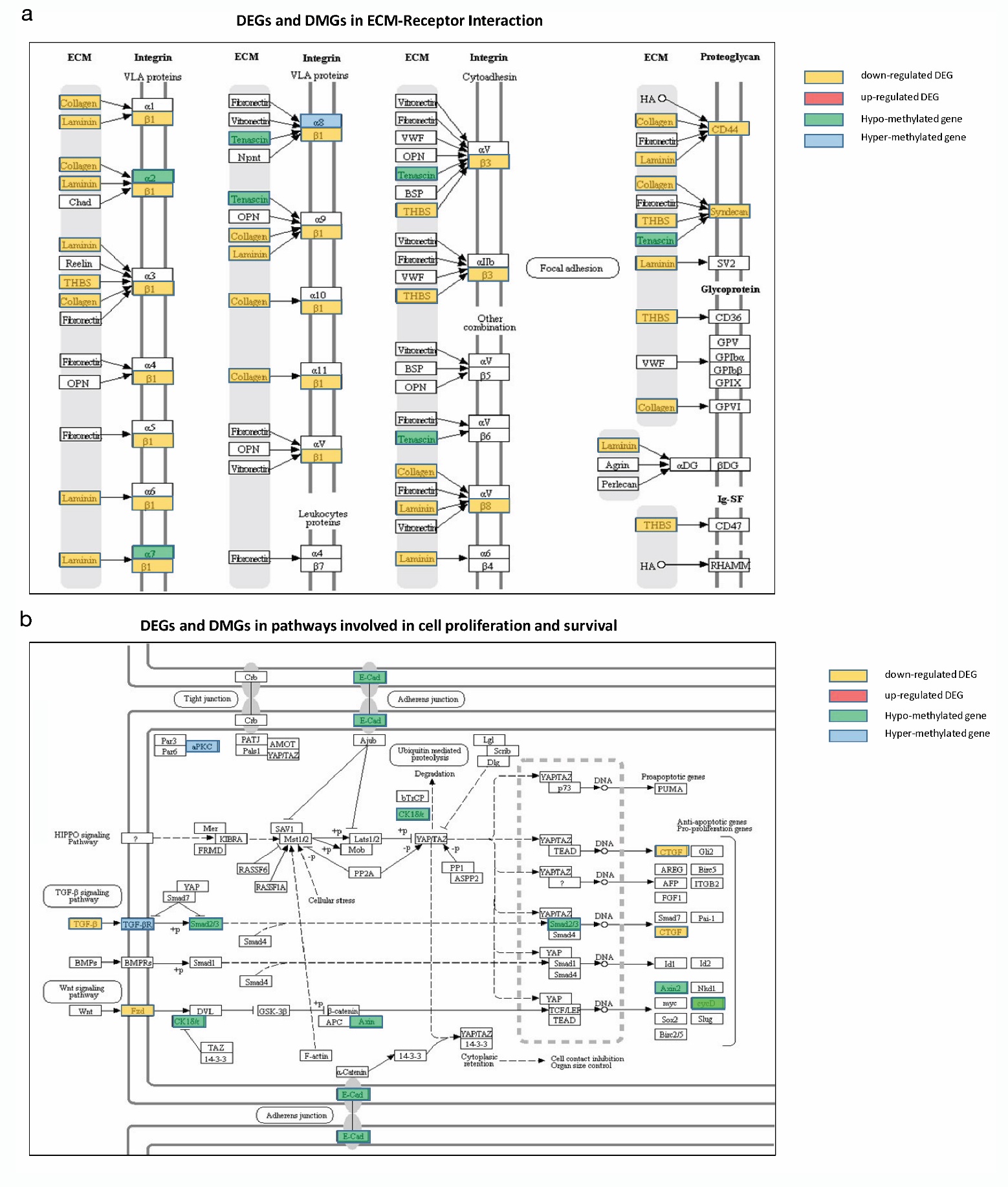

**Supplementary Figure 3. Integrated function analysis of the spaceflight-induced DMGs and DEGs found in EMC receptor, cell proliferation and survival**. (**a**) DEGs and DMGs found in Extracellular Matrix (ECM)-receptor interaction using KEGGs pathway analysis; (**b**) DEGs and DMGs found in cell proliferation and survival pathways using the DAVID enrichment analysis. The expression or methylation status of DEGs/DMGs was color-labelled as depicted by legends.
